# A general approach for identifying protein epitopes targeted by antibody repertoires using whole proteomes

**DOI:** 10.1101/641787

**Authors:** Michael L. Paull, Tim Johnston, Kelly N. Ibsen, Joel D. Bozekowski, Patrick S. Daugherty

**Affiliations:** Department of Chemical Engineering, University of California Santa Barbara CA, 93106, USA

## Abstract

Antibodies are essential to functional immunity, yet the epitopes targeted by antibody repertoires remain largely uncharacterized. To aid in characterization, we developed a generalizable strategy to identify antibody-binding epitopes within individual proteins and entire proteomes. Specifically, we selected antibody-binding peptides for 273 distinct sera out of a random library and identified the peptides using next-generation sequencing. To identify antibody-binding epitopes and the antigens from which these epitopes were derived, we tiled the sequences of candidate antigens into short overlapping subsequences of length k (k-mers). We used the enrichment over background of these k-mers in the antibody-binding peptide dataset to identify antibody-binding epitopes. As a positive control, we used this approach, termed K-mer Tiling of Protein Epitopes (K-TOPE), to identify epitopes targeted by monoclonal and polyclonal antibodies of well-characterized specificity, accurately recovering their known epitopes. K-TOPE characterized a commonly targeted antigen from *Rhinovirus A*, identifying three epitopes recognized by antibodies present in 83% of sera (n = 250). An analysis of 2,908 proteins from 400 viral taxa that infect humans revealed seven enterovirus epitopes and five Epstein-Barr virus epitopes recognized by >30% of specimens. Analysis of *Staphylococcus* and *Streptococcus* proteomes similarly revealed six epitopes recognized by >40% of specimens. These common viral and bacterial epitopes exhibited excellent agreement with previously mapped epitopes. Additionally, we identified 30 HSV2-specific epitopes that were 100% specific against HSV1 in novel and previously reported antigens. The K-TOPE approach thus provides a powerful new tool to elucidate the organisms, antigens, and epitopes targeted by human antibody repertoires.

## Introduction

Immunological memory allows for rapid antibody responses towards diverse antigens long after initial exposure. For example, the adaptive immune response to many vaccinations is often sustained throughout an individual’s lifetime [1]. This immunological information is archived within the genes encoding B-cell and T-cell receptors along with the corresponding receptor structures, but has proven difficult to characterize in a comprehensive manner. The ability to more fully interrogate immunological memory could reveal exposures to pathogens, commensal organisms, and allergens. Such information has proven useful for correlating antibody responses with disease outcomes to design more effective vaccines [2]. A detailed record of immune exposures can also facilitate the identification of biomarkers to diagnose infectious [3], autoimmune [4], and allergic conditions [5]. Furthermore, the capability to broadly characterize antibody repertoires at the epitope level could be used to identify conserved pathogen epitopes [6] and tumor specific antigen epitopes [7] to aid in therapeutic discovery.

A disease with prominent antibody responses is the common viral infection HSV, which causes human infections in the orofacial region (“cold sores”) and the genital region (“genital ulcers”) [8]. In 2012, the global prevalence of HSV1 was 3.7 billion people ages 0-49 [9] and the global prevalence of HSV2 was 417 million people ages 15-49 [10]. Diagnostic discovery generally focuses on diagnosing HSV2, since HSV2 infections can exacerbate HIV infections [10]. However, HSV1 and HSV2 contain the same genes [11] and the protein-coding regions of the HSV1 and HSV2 genomes share 83% sequence homology [12]. Therefore, researchers have often analyzed HSV glycoprotein G, since it differs substantially between the two HSV species [13]. In general, efforts have been limited to analyses of the surface-exposed envelope glycoproteins [14–17], using approaches such as microarrays [18]. Therefore, it would be novel to probe immunological memory using the entire proteomes of HSV1 and HSV2.

Immunological memory has been investigated extensively through sequencing the variable regions of B- and T-cell receptor encoding genes amplified from circulating cells [19]. These methods have proven useful for identifying receptor-encoding genes that associate with vaccination [20]. Nevertheless, such genetic information has not generally provided insight into the specific environmental antigens and epitopes targeted, unless they are known *a priori*. Furthermore, these methods require large specimen volumes (>10 mL) to obtain a sufficient quantity of cells [20]. Thus, there remains a need for methods that identify the diverse antigen targets of adaptive immunity.

Several methods have been developed to profile the protein epitopes of the secreted antibody repertoire [21]. Approaches have often focused on linear epitopes since 85% of epitopes contain at least one contiguous stretch of five amino acids [22]. By analyzing linear epitopes, researchers have identified sensitive and specific diagnostic epitopes for numerous diseases [21]. One common approach to epitope mapping is to generate short overlapping peptides by tiling candidate antigens. These peptides are then assayed for serum antibody reactivity in peptide microarray [23] or bacteriophage display library [24] formats. However, because these methods are biased towards specific organisms, they do not enable comprehensive or hypothesis-free immune evaluation. One strategy to overcome the limitations of tiling experiments is to use fully random peptide libraries [5,25,26]. Here, experiments are less biased and methods can analyze epitopes corresponding to a variety of organisms and antigens. A disadvantage of microarrays is that they are typically several orders of magnitude less diverse than peptide display libraries (e.g. 10^5^ [25] versus 10^10^ [5]), limiting the effectiveness with which current methods can achieve epitope discovery for low titer antibodies. In random library experiments, epitopes are typically discovered using *de novo* motif discovery by unsupervised clustering [27]. The most widely used algorithm for this purpose, MEME, scales approximately quadratically with the number of input sequences, making it less useful for analyzing large datasets resulting from next generation sequencing (NGS). While full-length antibody-binding peptides can be analyzed, the majority of the binding energy is typically derived from just 5-6 amino acids [28], thus other amino acids within the peptide will contribute noise. To rectify this problem researchers developed the IMUNE algorithm to reduce peptide datasets into statistically enriched patterns and cluster these patterns to build motifs [29].

A significant challenge for epitope mapping approaches is the association of epitopes and motifs with their corresponding antigens. Neither MEME nor IMUNE have the integrated capability to connect motifs to plausible antigens. Also, motifs identified through these methods often fail to reach the seven amino acids requirement for unambiguous identification of antigens within the full database of protein sequences [30]. Fundamentally, linear stretches in epitopes are typically less than seven amino acids in length [22], therefore, protein database searches of individual epitopes (such as through BLAST [31]) often fail to achieve statistical significance. Using multiple epitope matches within a single candidate antigen can increase the confidence of antigen prediction [26,32]. However, this method is insufficient for antigens with a single important epitope. Additionally, protein database searches are conducted using short amino acid sequences, therefore these searches do not fully leverage large quantitative binding datasets. To address these challenges, we present a general approach for associating epitopes with antigens using large peptide datasets. The K-mer Tiling of Protein Epitopes (K-TOPE) algorithm identifies epitopes by computationally tiling candidate antigens into k-mers, which are then evaluated within large datasets of antibody-binding peptides. Here, we demonstrate the utility of this approach by identifying linear epitopes within the proteomes of several prevalent infectious pathogens.

## Results

To enable the identification of protein epitopes bound by serum antibodies, we developed a method that uses a database of antibody-binding peptides to identify epitopes in known protein sequences (Fig 1). First, we selected peptides binding to an individual antibody repertoire within a specimen (serum or plasma) from a bacterial display peptide library with 10^10^ random 12-mer members. Then, we identified antibody-binding peptide sequences using NGS. To allow for the manipulation of 20^5^ (3.2 million) k-mers rather than full-length peptides, we processed peptides into subsequences and evaluated the enrichments of all k-mers of length 5 [29]. Next, K-TOPE tiled candidate antigen sequences, such as from a proteome, into overlapping k-mers. K-TOPE used the enrichment values for these k-mers to construct an enrichment histogram across the length of each protein sequence. The frequency value at each sequence position in the histogram was proportional to the enrichment of k-mers that included that position. Specifically, for all k-mers overlapping a position, we summed the log base 2 of the k-mer enrichment. Thus, higher frequency values at a position in a protein sequence corresponded to a greater probability that a position was included in an epitope. Epitopes were extracted from the maxima in the histogram and scored based on their area under the curve (AUC). Finally, epitopes were assigned an “epitope percentile” based on their rank in a list of scores generated from random proteins.

**Fig 1.**
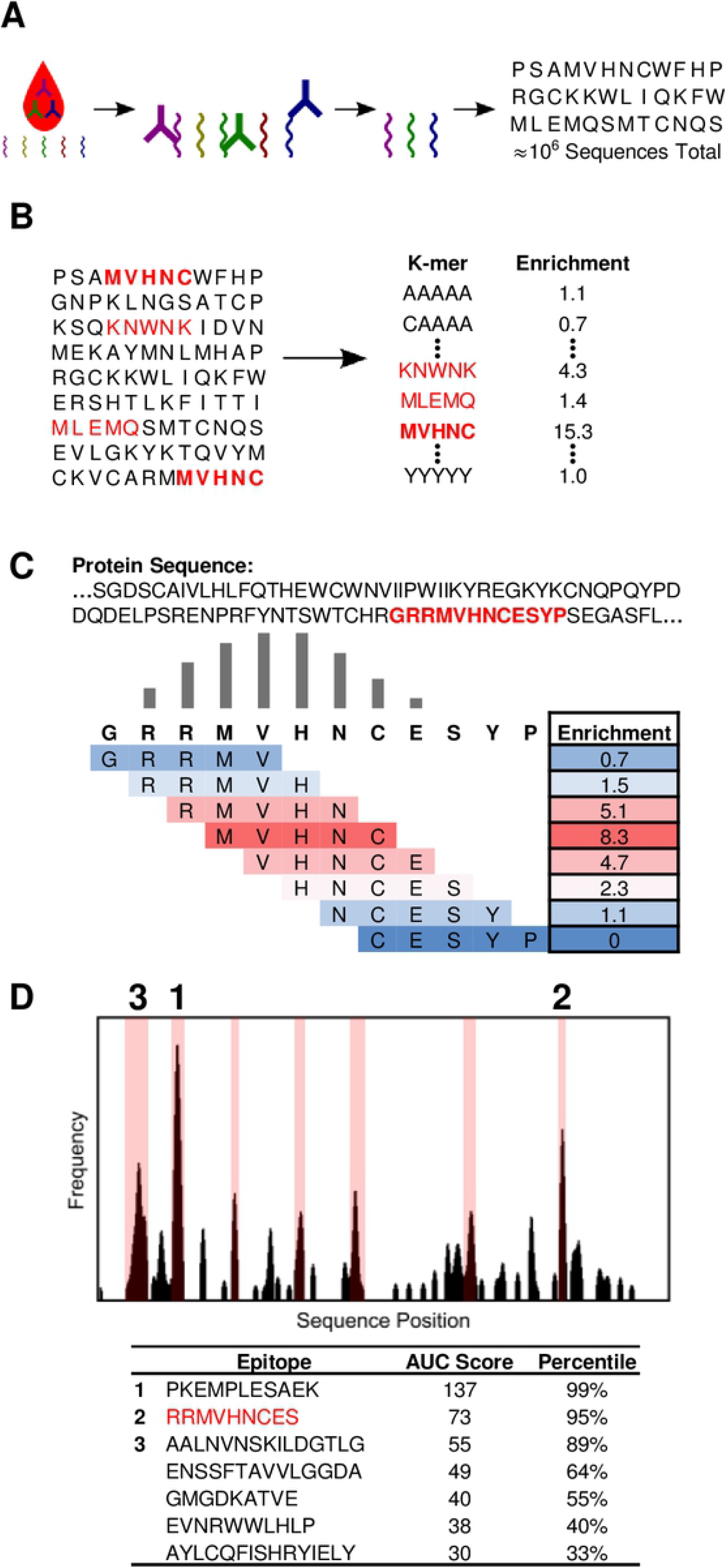
K-TOPE determines epitopes by tiling proteins into k-mers. (A) The input to the algorithm is a dataset of approximately 10^6^ peptides that were bound by serum antibodies. (B) All 5-mers are evaluated for their enrichment in the list of peptides. (C) A portion of a protein sequence is tiled into 5-mers which are weighted by their enrichment. This determines a “frequency” value for each position in the sequence. (D) The frequency value for each position in a protein sequence is plotted as a histogram. Possible epitopes are highlighted in pink on the graph. Epitope sequences, area under the curve (AUC) scores, and significance percentiles are displayed.

To assess the utility of K-TOPE, we first determined epitopes for monoclonal and polyclonal antibodies that bind specific, well-defined epitopes in cMyc, V5, and amyloid beta. We spiked these antibodies into serum at a final concentration of 25 nM and then selected and identified binding peptides. K-TOPE identified epitopes that corresponded closely to the previously reported epitopes of these antibodies (Fig 2). Importantly, the enrichment histograms generated by antibodies spiked into background serum or buffer were nearly identical (S1 Fig), suggesting that the noisy serum environment minimally affected epitope identification.

**Fig 2.**
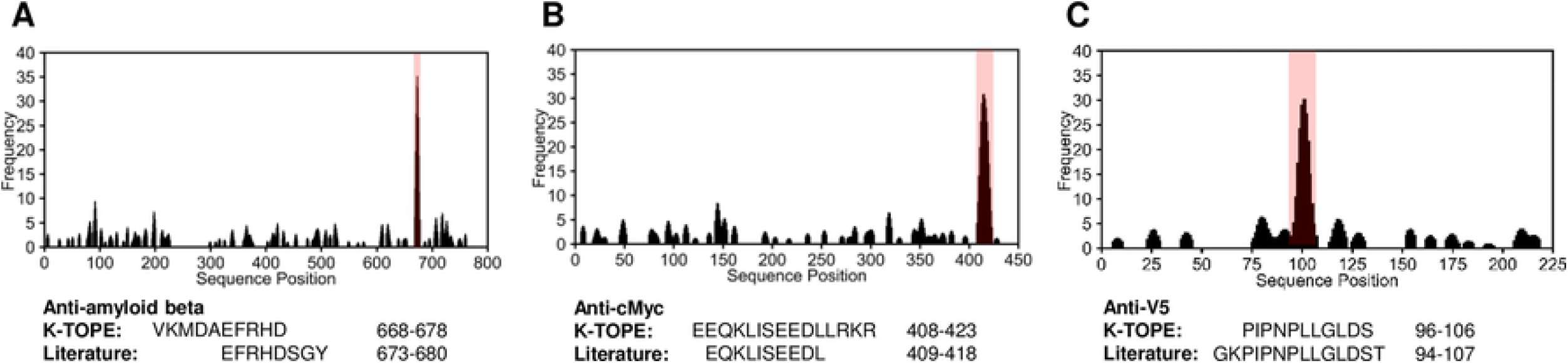
K-TOPE found epitopes for antibodies with known specificity spiked into serum. Histograms for antibodies with known specificity against amyloid beta (P05067), cMyc (P01106), and V5 (P11207) had prominent epitopes (in pink). (A) K-TOPE analysis of amyloid beta determined the epitope VKMDAEFRHD (668-678). This antibody was raised to whole protein and is known from literature to have a conformation-specific discontinuous epitope that maps to segments EFRHDSGY (673-680) and ED (692-693). (B) K-TOPE analysis of cMyc determined the epitope EEQKLISEEDLLRKR (408-422). This antibody was raised to AEEQKLISEEDLLRKRRE (407-424). (C) K-TOPE analysis of V5 determined the epitope PIPNPLLGLDS (96-106). The antibody was raised to GKPIPNPLLGLDST (94-107).

To identify “public epitopes” conserved across many individuals, epitopes were generated for each specimen individually and then clustered. Although many private epitopes were identified for each specimen in this process, we focused on the far smaller set of public epitopes to facilitate comparison with previous literature. Given the ubiquity of exposure to the upper respiratory pathogen *Rhinovirus A*, we validated the approach by identifying epitopes within its genome polyprotein. Using a unique set of 250 serum specimens, we identified epitopes within *Rhinovirus A* that were targeted by 30% or more of the specimens (Fig 3A). Of the 250 specimens, 87% exhibited binding to at least one of these consensus epitopes (Fig 3B). Three of these epitopes were located within positions 570-620 (Fig 3C), in the antigenic attachment region of VP1. A fourth epitope within the VP2 region of the *Rhinovirus A* genome polyprotein was targeted by 43% of the population.

**Fig 3.**
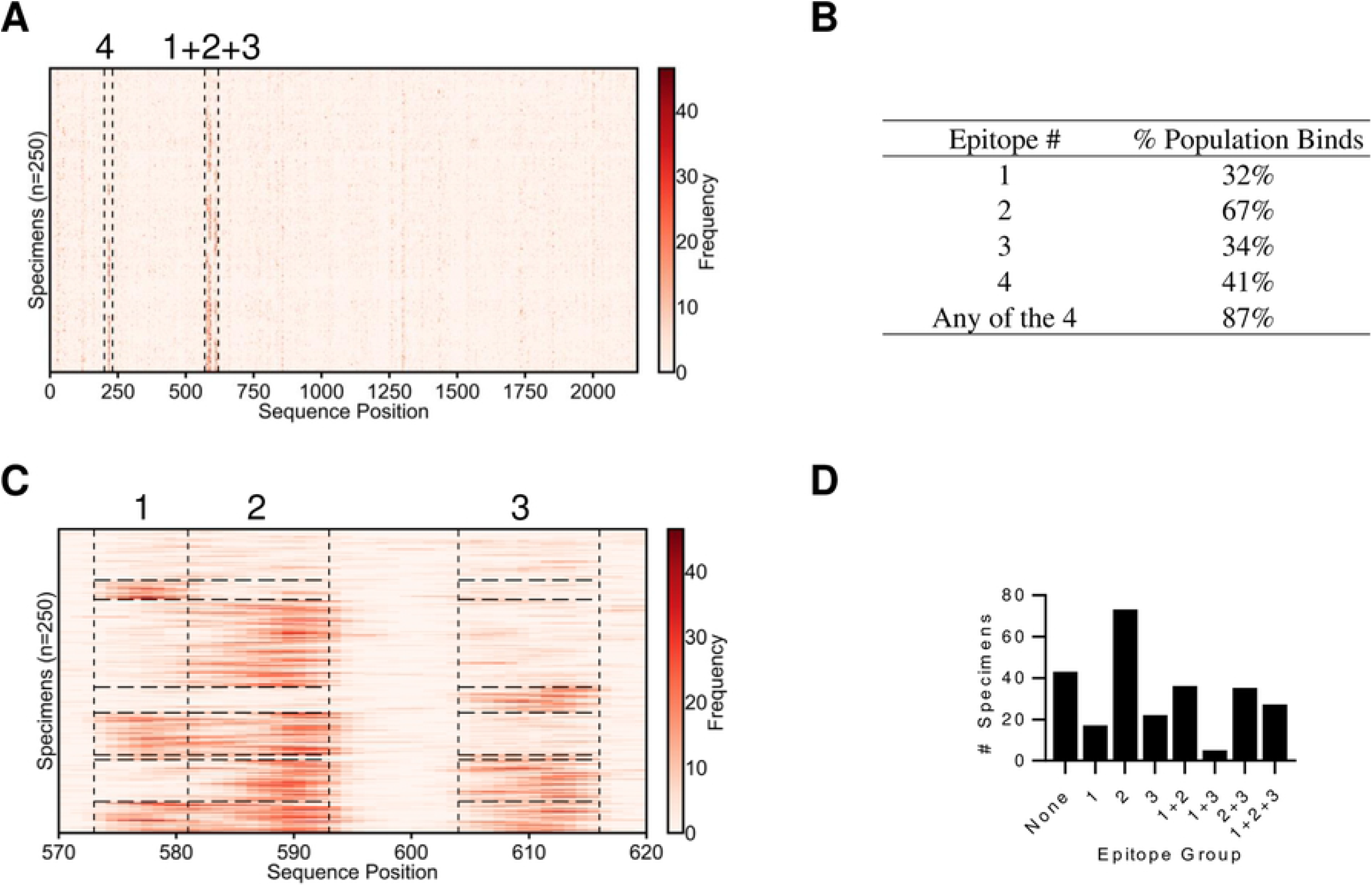
K-TOPE identified four epitopes in the *Rhinovirus A* genome polyprotein. (A) K-TOPE was applied to the *Rhinovirus A* genome polyprotein (P07210) for 250 specimens. Histograms for all specimens are shown as rows in a heat map. The specimens have been clustered such that specimens that bind the same epitopes are adjacent. Regions that contain epitopes are outlined by dotted lines. (B) A table of the percentage of the population that bound each epitope. For instance, Epitope 1 is the percentage of specimens that targeted “1”, “1+2”, “1+3”, “1+4”, “1+2+3”, “1+2+4”, “1+2+3+4”. (C) The region from positions 570-620 is divided into 3 sections that correspond to distinct epitopes. These epitopes are consensus epitopes which were present in >30% of the 250 specimens. (D) Bar graph showing membership in different epitope groups. For example, a specimen that binds epitopes 2 and 3 will belong to epitope group “2+3”. In this population, 87% of the specimens bound at least one of the consensus epitopes. The sequences of the epitopes were 1: QNPVENYI, 2: DSVLEVLVVPN, 3: APALDAAETGHT, and 4: NHTHPGEQG.

To assess trends in the population, each specimen was assigned into one of eight groups based on which of the three VP1 epitopes were bound (Fig 3D). Notably, epitope binding was not independent, since the group of specimens targeting all three epitopes was 44% larger than expected and the group targeting epitopes ‘1+3’ was 50% smaller than expected (S1 Table). The average age of the subset of specimens of known age (n=138) was 35 years, however, the epitope group targeting all three epitopes had an average age of 17, and the epitope group targeting none of the epitopes had an average age of 50 (S2 Table). Thus, people who targeted fewer *Rhinovirus A* epitopes tended to be older.

Next, we investigated the utility of using K-TOPE to identify epitopes within a set of 2,908 proteins from 400 viral taxa with human tropism. This approach yielded 29 epitopes that were bound by at least 30% of all specimens (Table 1). The prevalence of each epitope is noted, which is defined as the proportion of specimens that bound the epitope. Some of these epitopes have been reported previously [6,33–35]. Thus, a modest number of prominent linear viral epitopes were bound by >30% of the specimens analyzed. A common antigen identified from this analysis was Epstein-Barr nuclear antigen 1 (EBNA1) from Epstein-Barr virus (EBV), which is expressed in EBV-infected cells [36]. Additionally, the epitopes identified for the enterovirus genus were consistent with the epitopes identified for *Rhinovirus* A, which is a species in that genus (Fig 3). Several of the epitopes were likely due to false discovery (e.g., Mayaro virus and Lyssavirus), since these viruses are uncommon in a general population. There is an intrinsic lower limit on false positives since antibodies only bind 5-6 amino acids, which is not enough information to uniquely specify a protein subsequence. This limitation is especially pronounced among evolutionarily related proteins in closely related species.

**Table 1.**
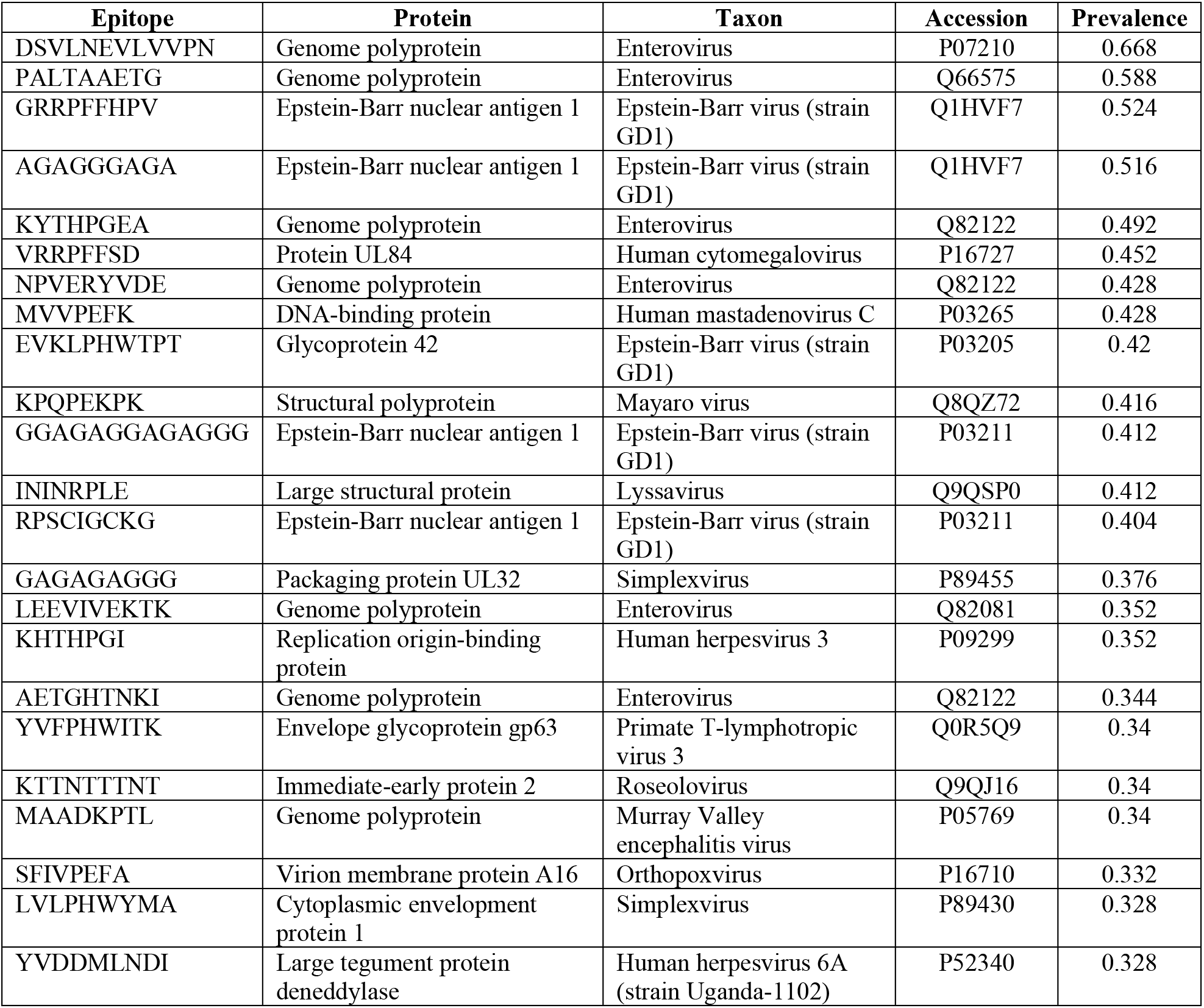
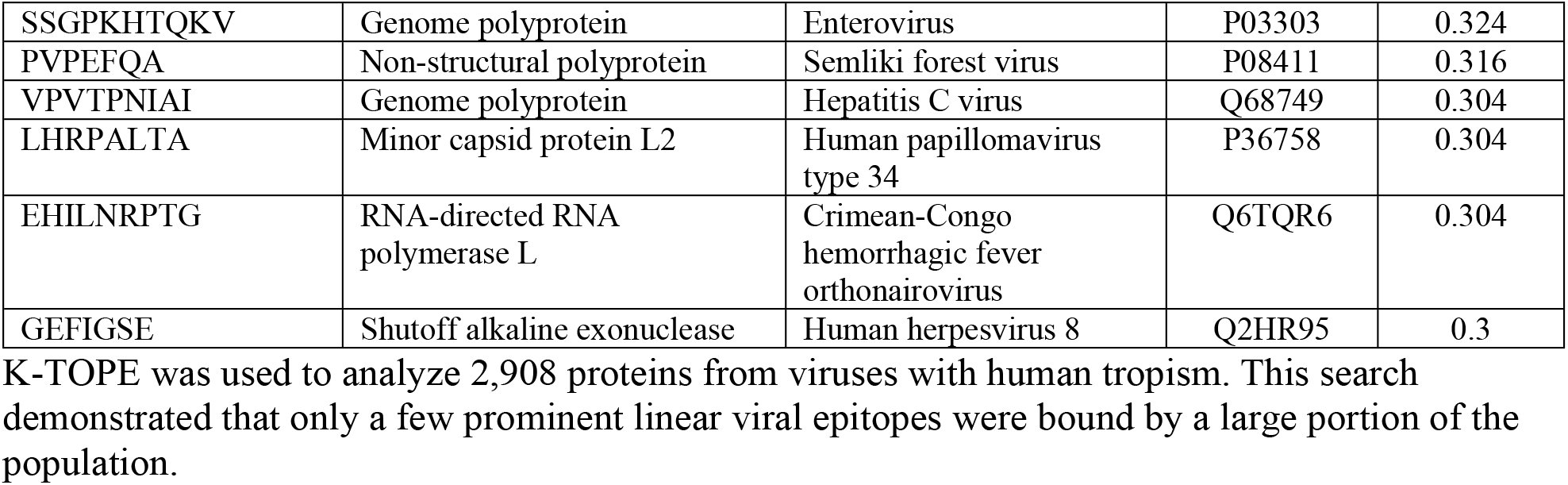
A collection of 29 viral epitopes to which >30% of 250 specimens bound.

We performed a similar analysis for the proteomes of the genera *Streptococcus* and *Staphylococcus*, which are common bacterial human pathogens with 2,976 and 3,071 proteins in their respective proteomes. K-TOPE was used with each of these proteomes to determine epitopes bound by >30% of a population of 250 specimens, yielding 9 epitopes for *Streptococcus* and 13 epitopes for *Staphylococcus* (Table 2). The epitope LIPEFIG(R) in ATP-dependent Clp protease ATP-binding subunit ClpX was the most prevalent *Streptococcus* epitope and second most prevalent *Staphylococcus* epitope. Therefore, K-TOPE could not determine which genus generated this epitope. The most prevalent *Staphylococcus* epitope was PTHYVPEFKGS from extracellular matrix protein-binding protein emp, which is a known virulence factor [37]. For *Streptococcus*, the second most prevalent epitope was GQKMDDMLNS from the highly antigenic Streptolysin O protein [38]. This epitope falls within a 70 amino acid range in Streptolysin O that is known to bind antibodies [39]. The sequence “DKP” was present in 5/9 *Streptococcus* epitopes and the sequence “PEFXG” was present in 6/13 *Staphylococcus* epitopes (Table 2). Therefore, there are multiple candidate antigens that may correspond to these highly enriched sequences.

**Table 2.**
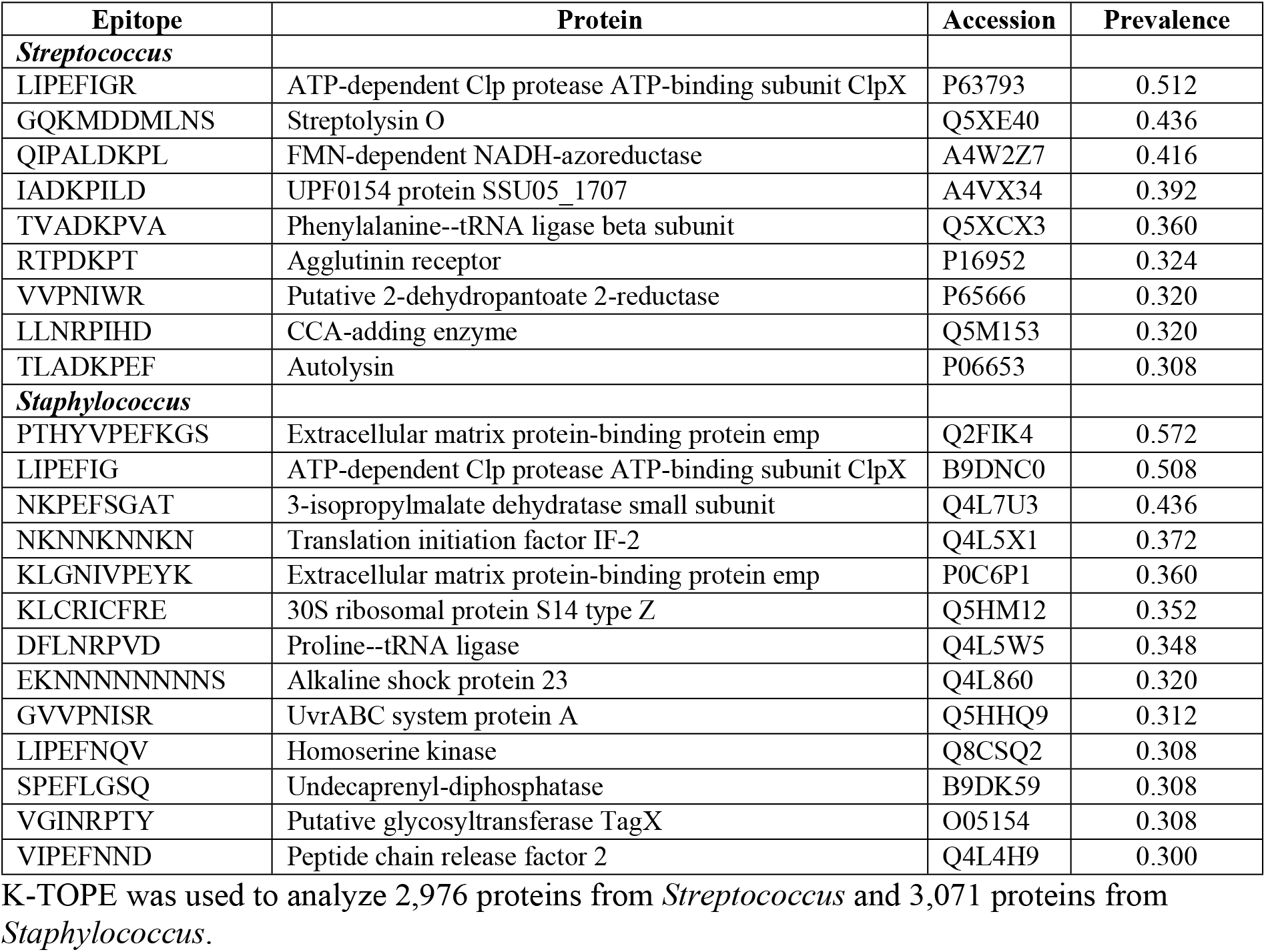
Epitopes in the proteomes of the genera *Staphylococcus* and *Streptococcus* which were bound by >30% of 250 specimens.

The most prevalent epitopes identified through proteome searches were validated by comparison to previously reported epitopes. We chose to analyze the viral proteins EBNA1 from EBV and the *Poliovirus 1* genome polyprotein (representing Enterovirus), which were present five and seven times, respectively, in Table 1. Bacterial proteins chosen for validation were Streptolysin O, corresponding to the second most prevalent *Streptococcus* epitope (Table 2), and Extracellular matrix protein-binding protein emp, corresponding to most prevalent *Staphylococcus* epitope (Table 2). In all cases, K-TOPE found prominent peaks in the histograms that corresponded to reported epitopes (Fig 4) [6,33,35,40]. Additionally, K-TOPE identified an immunogenic region of GA-repeats from positions 100-350 in the analysis of EBNA1 [23]. We used a nonparametric statistical test to assign significance to the overlap between K-TOPE epitopes and known epitopes. Using this method, all epitopes evaluated using K-TOPE had P-values below 0.05 (Fig 4C).

**Fig 4.**
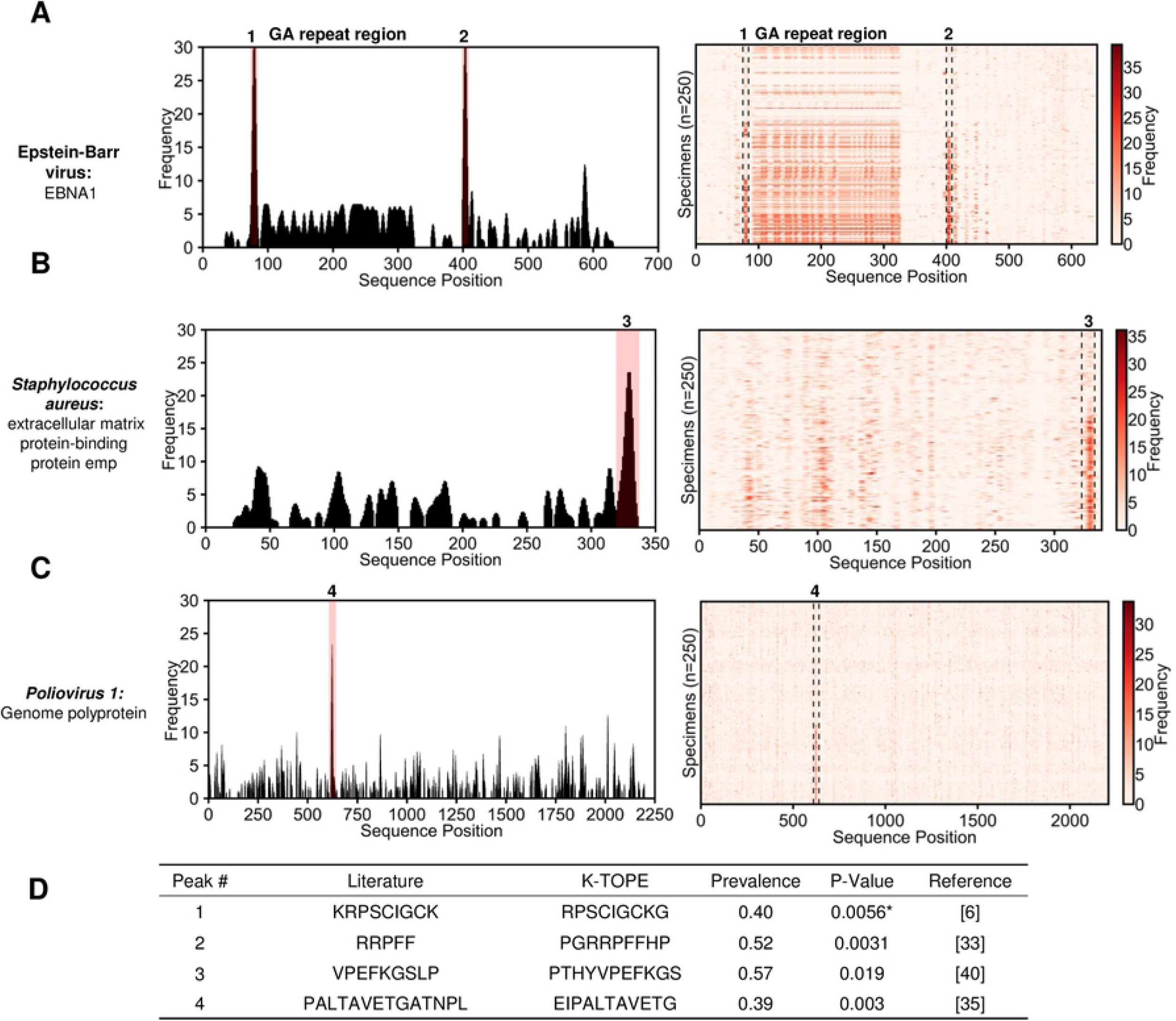
Epitopes identified through proteome searches were validated using literature-reported epitopes. In (A), (B), and (C), a histogram is shown for a single specimen with significant peaks (in pink). To the right of the histogram is a heat map for 250 specimens. For (A), there is a region of antigenic GA-repeats from positions 100-350. The table in (D) provides the statistical significance of agreement between literature epitopes and K-TOPE epitopes for the labeled peaks in (A), (B), and (C). The UniProt accessions used for this analysis were P03211 for EBNA1, Q8NXI8 for extracellular matrix protein-binding protein emp, and P03300 for *Poliovirus 1* Genome Polyprotein. Statistical tests where epitopes with >50% GA content were removed are denoted by an asterisk “*”. All identified epitopes had p-values below 0.05.

To identify HSV species-specific epitopes, we analyzed 12 HSV2 specimens and 10 HSV1 specimens. Since these viruses share many of the same proteins in their proteomes [11], HSV1 specimens were appropriate controls for HSV2 specimens and vice-versa. To begin, we identified species-specific epitopes in glycoprotein G, which is a protein that varies significantly between the two species (Fig 5) [41]. There was a single HSV1 epitope, PMPSIGLEE, bound by 40% of HSV1 specimens and a single HSV2 epitope, GGPEEFEGAGD, bound by all HSV2 specimens. This HSV2-specific epitope aligned well with previous epitopes found for glycoprotein G2 [13,42,43] (Table 3). Also, this epitope has been validated as an HSV2-specific diagnostic [44,45]. The HSV1-specific epitope was also similar to the previously reported epitope DHTPPMPSIGLE [18]. Interestingly, the two HSV-specific epitopes terminated in an identical 7-mer sequence EGAGDGE (PMPSIGLEEEEEE**EGAGDGE** and GGPEEF**EGAGDGE**) [42]. This suggests that the regions containing these epitopes may be evolutionarily or structurally related targets of the immune system.

**Table 3.**
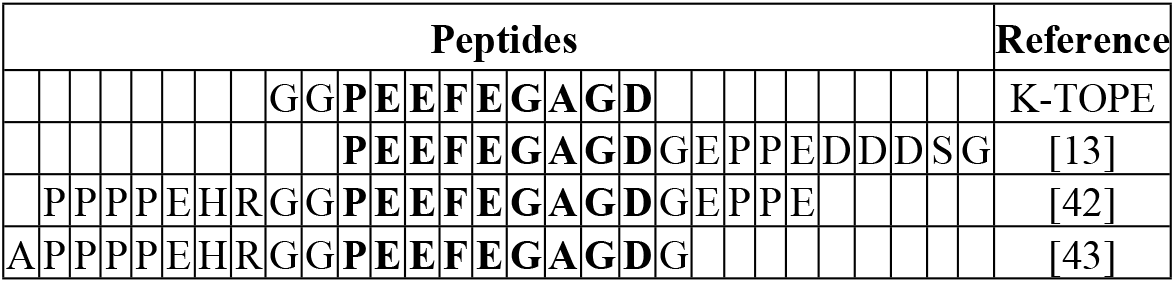
Alignment of an HSV2-specific glycoprotein G2 epitope with previously reported epitopes.

**Fig 5.**
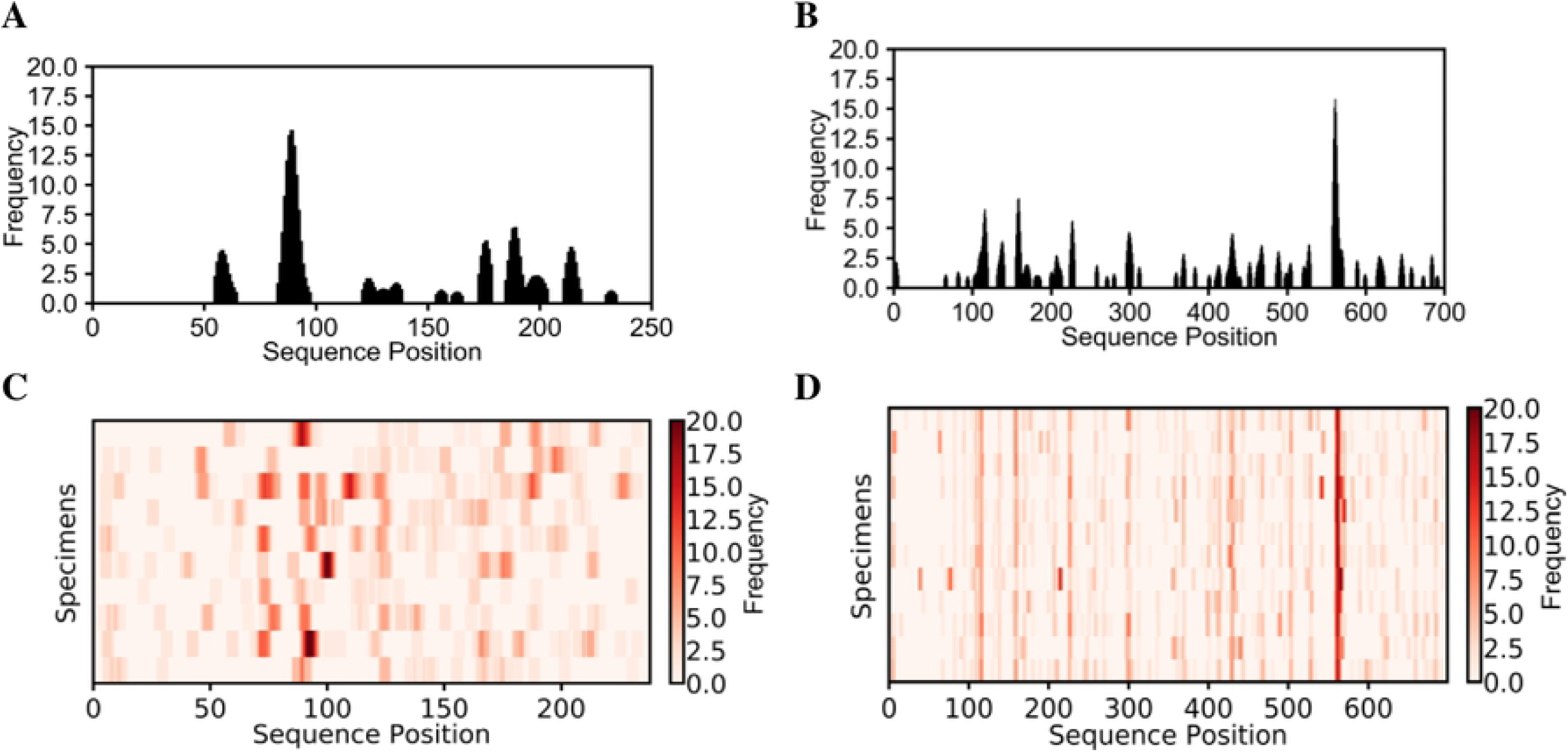
K-TOPE identified epitopes for glycoprotein G1 using HSV1 specimens and for glycoprotein G2 using HSV2 specimens. For glycoprotein G1, a representative histogram for a single specimen is shown in (A) and a heat map for all HSV1 specimens is shown in (C). For glycoprotein G2, a representative histogram for a single specimen is shown in (B) and a heat map for all HSV2 specimens is shown in (D). There was a single epitope identified for each protein.

To identify candidate HSV species-specific epitopes, we analyzed the HSV1 and HSV2 proteomes. We identified 30 HSV2-specific epitopes that were 100% specific with prevalence > 30% (Table 4). Notably, 11 of these epitopes were bound by all HSV2 specimens. K-TOPE identified a glycoprotein C epitope PRTTPTPPQ with 83% prevalence which was contained in a previously identified epitope RNASA**PRTTPTPPQ**PRKATK [18]. In contrast to the numerous HSV2-specific epitopes, only 4 HSV1-specific epitopes were identified, and the highest prevalence achieved was only 40% (Table 5). One of these epitopes, RIRLPHI, overlapped with the previously identified epitope HRRTRKAPK**RIRLPHI**R [46] in the well-described antigen glycoprotein D [17]. One possible explanation for the discovery of fewer HSV1-specific epitopes is that the HSV2 specimens had high IgM levels, whereas the HSV1 specimens had high IgG levels. Since high IgM levels occur with severe recurrent herpes infections [47], we would expect the high IgM HSV2 sera to yield more epitopes.

**Table 4.**
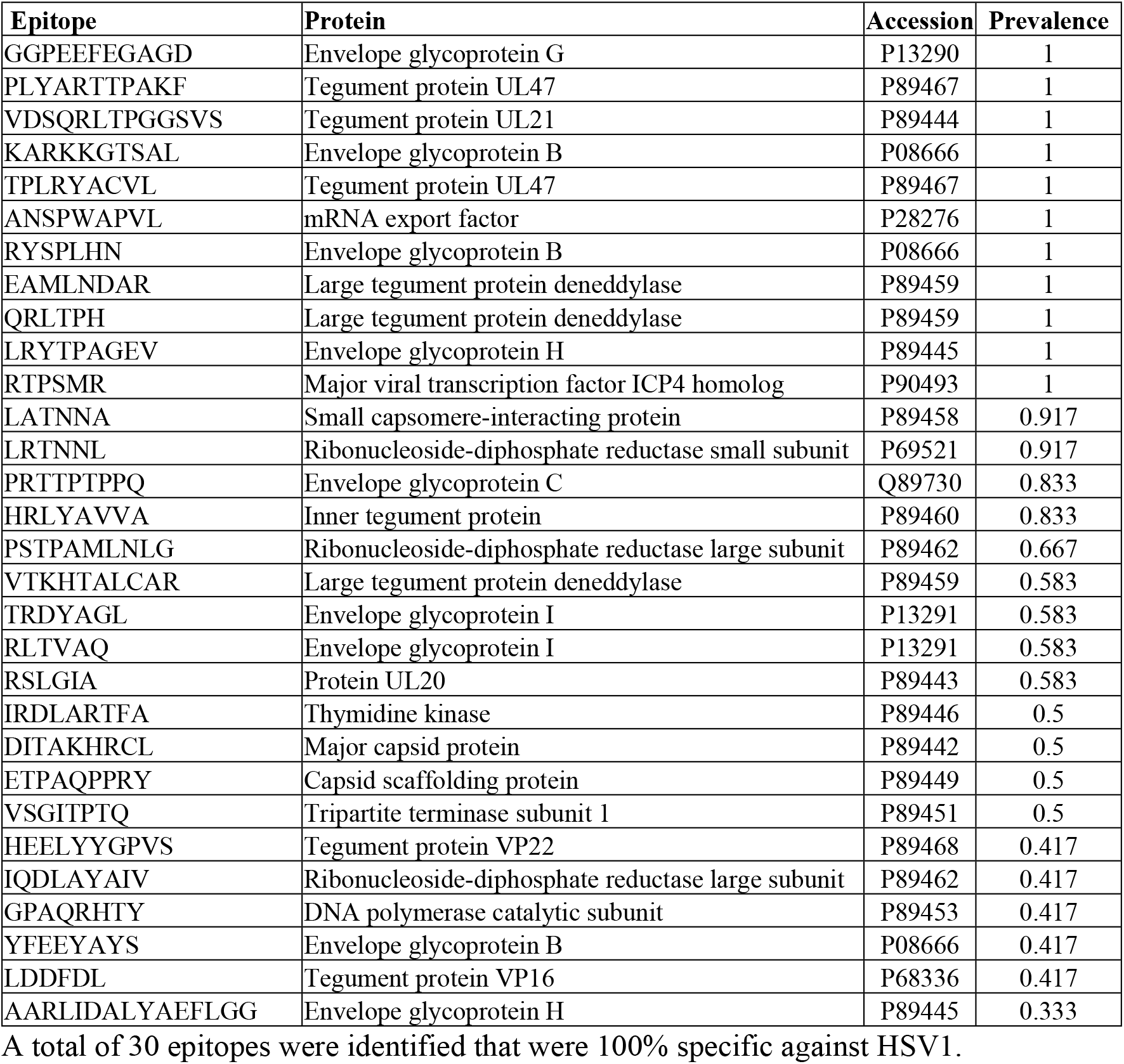
HSV2-specific epitopes were identified.

**Table 5.**
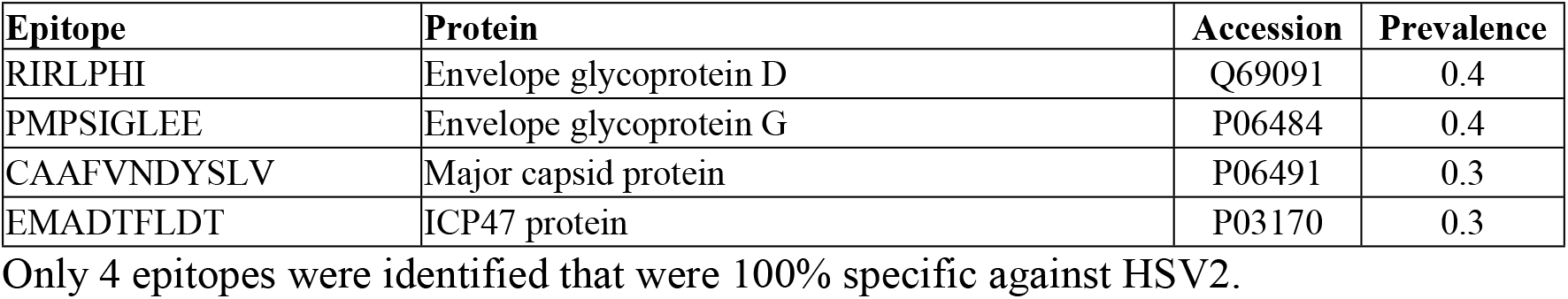
HSV1-specific epitopes were identified.

| Epitope | Protein | Accession | Prevalence |
| --- | --- | --- | --- |
| RIRLPHI | Envelope glycoprotein D | Q69091 | 0.4 |
| PMPSIGLEE | Envelope glycoprotein G | P06484 | 0.4 |
| CAAFVNDYSLV | Major capsid protein | P06491 | 0.3 |
| EMADTFLDT | ICP47 protein | P03170 | 0.3 |

Only 4 epitopes were identified that were 100% specific against HSV2.

We sought to determine whether the HSV2-specific epitopes were contained in proteins that differed between the HSV species [41]. We determined 8 HSV2-specific epitopes with sequences that were contained in both HSV proteomes (S3 Table). Our analysis suggested that these epitopes were only targeted by HSV2 specimens, despite their presence in the HSV1 proteome. Thus, even sequences that are conserved between species could serve as species-specific targets.

## Discussion

Here, we present a generalizable methodology for identifying epitopes within candidate immunogenic proteins. By tiling proteins into k-mers and evaluating those k-mers in a database of antibody-binding peptides, we determined epitopes for individuals and a population. Importantly, we have demonstrated that K-TOPE can identify disease-specific epitopes and antigens. One of the main features of this approach is that it combines k-mers to determine composite epitopes that may not explicitly exist in the peptide dataset. Another important element is using an antigen sequence to identify epitopes, thereby surmounting the 7 amino acid requirement for successful antigen identification [30].

The K-TOPE approach to epitope mapping differs from reported methods in several important ways. While proteome-derived peptide libraries have been used to identify disease-specific epitopes [33,48], these methods lack the flexibility to examine multiple proteomes. For instance, separate libraries would be required to analyze both HSV1 and HSV2. Even a library that contains peptides spanning all viral proteomes cannot easily be extended to much larger bacterial or parasitic proteomes [24]. A disadvantage of microarrays is that they have far lower 5-mer coverage (~27% [32]), than surface display (~100%) which could limit the application of k-mer approaches. Other algorithms have been developed that identify binding motifs in peptide datasets, but they lack the integrated capability to connect motifs to protein antigens [49,50]. Also, the direct method of aligning peptides to sequences becomes computationally infeasible with a large number of peptides and candidate antigens [51].

The heterogeneity of experimental approaches complicates the validation of putative epitopes and their associated antigens. The Immune Epitope Database (IEDB) has an all-inclusive representation of information [52], which may not reflect important distinctions in experimental platforms, specimens, and data analysis techniques. For instance, there are likely numerous false positive epitopes for highly studied organisms and few identified epitopes for poorly studied organisms. Also, there is a lack of quantitative data reported for epitopes [53], such as the proportion of a given population that binds an epitope. To address this lack of information, we first used K-TOPE to analyze specimens for responses to common pathogens in a general population. This allows newly identified “public epitopes” to be benchmarked by nearly any set of serum specimens. We required that a proportion of the population bind an epitope to reduce false positives. Although analysis of the variation in private epitopes could be valuable for understanding the variation in immune responses, it would complicate validation. We determined public epitopes in *Rhinovirus A* and showed that people who targeted fewer *Rhinovirus A* epitopes tended to be older, perhaps due to immunosenescence [54], reduced pathogen exposure, or a lower incidence of rhinovirus infections [55]. With a diverse group of specimens, it was possible to confirm that the RRPFF epitope in EBV’s protein EBNA1 is a very commonly targeted epitope [33]. Since the specimens used to determine public epitopes were not assayed for responses to pathogens, acute and chronic infections could not be readily distinguished from prior infections. These public epitopes could be further validated using specimens with acute infections or using longitudinal studies to determine if these epitopes appear upon vaccination [56]. We did not find epitopes corresponding to measles or rubella vaccination, which is consistent with a recent study that comprehensively identified viral epitopes [57]. This implies that for these viruses, high titer antibodies targeting linear epitopes may not be present. For HSV1 and HSV2, we determined whether an epitope was specific by analyzing specimens infected by both virus species. Unexpectedly, we demonstrated that even epitopes present in the conserved regions of both species’ proteomes could be species-specific. The difference in binding was likely due to differences in the structure and post-translational modifications of the proteins. For the HSV analysis, we validated epitopes using previous studies, however, it was difficult to know *a priori* whether a non-validated epitope was novel or spurious. In general, since studies use different specimens, experiments, and computational analyses, it is unlikely for the epitopes of two studies to completely coincide.

K-TOPE provides a new tool for identifying diagnostic biomarkers, vaccine components, and candidate therapeutic targets. This approach could be used in the iterative process of designing a vaccine, since it would be useful to know which epitopes are elicited in a population by vaccination. Vaccine formulation could be altered to maximize the percentage of the population that targets epitopes associated with a positive disease outcome [2]. K-TOPE could also enable the development of diagnostics that assign disease based on the presence of epitopes. Since this method only involves a single experimental screen, in principle multiple diseases could be simultaneously diagnosed [58]. By searching for consensus epitopes in a disease group that are absent in a control group, K-TOPE can discover disease-specific epitopes. For an autoimmune disease, the entire human proteome could be analyzed to determine autoantigen epitopes [33]. Similarly, using clinical histories of viral infection, K-TOPE can analyze the proteomes of suspected pathogens to link epitopes to infections [24]. With specimens that have HLA information, it could be possible to detect a correlation between HLA type and bound epitopes [59]. This could have implications for how we determine genetic predisposition to immunological disease.

There are important limitations to the conditions in which this approach could be successful. First, this approach is currently limited to the identification of linear epitopes. However, since 85% of epitopes have at least one linear stretch of five amino acids [22], conformational epitopes with linear segments may be represented in the datasets. We chose to focus on linear epitopes since methods that identify conformational epitopes often require 3D protein structures, which are scarce relative to the number of protein sequences. This report focuses on epitopes from common pathogens which are high-titer, but it could be difficult to detect rare antibody epitopes. Methods that selectively deplete out high-titer antibodies could prove effective for probing rare antibodies [60]. Another limitation is that protein sequences tend to have a large degree of conservation and redundancy [61], as demonstrated by the false positives found in the viral epitope search. Thus, even for analyses of non-immunogenic proteomes, false positives will occur due to evolutionary or coincidental sequence overlap with immunogenic proteomes. The issue of false positives can be partially allayed by deliberately choosing the set of investigated proteins, such that all proteins are plausible candidate antigens. Sequence conservation was demonstrated with the Enterovirus epitope PALTAVETGATNPL [35], as well as with the *Human herpesvirus 6A* epitope YVDDMLNDI (Table 1) which shares the k-mer “DDMLN” with the *Streptococcus* epitope GQKMDDMLNS (Table 2). Generally, if an epitope sequence is present identically in multiple antigens, all candidate antigens should be considered equally plausible without further biological, epidemiological, or experimental information. It is important to note that one of the purposes of K-TOPE is to reduce thousands of candidate proteins to a small set of proteins that can be experimentally validated.

In summary, the present approach enables the discovery of epitopes within the proteomes of any organism whose sequence is deposited into the protein database. The challenge of associating epitopes with antigens can be surmounted by transforming sets of antibody-binding peptides to k-mers and tiling proteins of interest. Advancements upon this paradigm may enable comprehensive immunological evaluations from serum and other biological tissues.

## Materials and methods

### Strains and reagents

*E. coli* strain MC1061 was used with surface display vector pB33eCPX for all library screening experiments. Protein A/G magnetic beads were from Thermo Scientific Pierce. Antibodies with known specificity included C3956 rabbit anti-c-Myc polyclonal antibody (Sigma), anti-beta amyloid 1-42 antibody [mOC31] - conformation-specific (ab201059) (Abcam), and rabbit V8137 Anti-V5 polyclonal antibody (Sigma). Antibodies were spiked into healthy donor serum at a concentration of 25 nM. All sera (n=273) were obtained as deidentified specimens from biobanks according to institutional guidelines, (Biosafety authorization numbers #201417, #201713), and handled according to CDC-recommended BSL2 guidelines.

### Bacterial peptide display and sequencing

The bacterial peptide display screening protocol was carried out as previously described [29,62]. Briefly, an *E. coli* library displaying approximately 8 billion different 12-mer peptides was combined with 1:100 diluted serum. We used magnetic selection with Protein A/G beads to isolate bacterial cells with bound antibodies. Then, we confirmed that this isolated fraction of bacteria bound antibodies using flow cytometry. Amplicons were prepared from the isolated fraction for sequencing using the Illumina NextSeq.

### Protein databases

Protein sequences were obtained from UniProt or by using the Biopython module [63]. Accessions for proteins are noted in figures and figure captions. For the epitope validation, accessions were chosen that reference the most highly annotated version of the proteins identified in Table 1 and Table 2. The list of random proteins used for statistical analysis was obtained through a UniProt search of “reviewed:yes”. The viral proteome search used a Uniref search of “uniprot:(host:"homo sapiens" reviewed:yes fragment:no) AND identity:0.9” and yielded 2,908 proteins. The *Staphylococcus* proteome search used a Uniref search of “uniprot:(taxonomy:“Staphylococcus [1279]” fragment:no reviewed:yes) AND identity:0.9” and yielded 3,071 proteins. The *Streptococcus* proteome search used a Uniref search of “uniprot:(taxonomy:“Streptococcus [1301]” fragment:no reviewed:yes) AND identity:0.9” and yielded 2,976 proteins. HSV analysis used a UniProt search of “reviewed:yes AND organism:“Human herpesvirus 1 (strain 17) (HHV-1) (Human herpes simplex virus 1) [10299]” AND proteome:up000009294” for HSV1, yielding 73 proteins and a Uniprot search of “reviewed:yes AND organism:“Human herpesvirus 2 (strain HG52) (HHV-2) (Human herpes simplex virus 2) [10315]” AND proteome:up000001874” for HSV2, yielding 72 proteins.

### Selection of literature epitopes

For EBNA1, RRPFF was chosen because it was noted that RRPFF antibodies were found in the serum of healthy individuals [33]. KRPSCIGCK was noted as an EBNA1 epitope that was preferentially targeted by pre-eclamptic women, but was also targeted by healthy controls [6]. The motif XPEFXGSXX was discovered and inferred to correspond to VPEFKGSLP in *Staphylococcus aureus* using protein database searches [40]. For *Poliovirus 1*, the epitope PALTAVETGATNPL was found to be a cross-reactive epitope in many enteroviruses [35].

### Sequence processing

All software files are posted on GitHub (https://github.com/mlpaull/KTOPE) and all 278 antibody-binding peptide files are available on Dryad (doi:10.5061/dryad.v7d0350). The imune-processor.jar file is available for research, non-profit, and non-commercial use and requires a license for commercial use. All other software is available under the MIT license. The algorithms for generating nonredundant sequence lists from FASTQ files, outputting enrichment values for subsequences, and exhaustively calculating k-mer statistics were adapted from IMUNE (imune-processor.jar and calculate-patterns.jar) [29]. We added the capability to start with lists of peptides rather than NGS data. The enrichment of a k-mer is defined as the ratio of the number of observations of the k-mer to the “expected” number of observations. The “expected” value is calculated as the product of the total number of sequences, the number of frames the k-mer could fit in the sequences, and the probability of the k-mer appearing based on amino acid usage. If a k-mer’s enrichment is above the “enrichment minimum” (2.0 for this study), it is used in K-TOPE. K-mers need to be calculated only once per specimen. All interaction with IMUNE-derived code is through a Python module which sets up a folder hierarchy and acts as a wrapper for IMUNE-derived code (imuneprocessor.py). These programs are memory and hard-drive intensive and it is recommended to have at least 16 GB of free RAM and 100 GB of hard-drive space. Analysis was carried out on a Dell Optiplex 9020 with an Intel^®^ Core™ i7-4790 CPU @ 3.60 GHz, 64-bit operating system, and 32.0 GB of RAM. Processing FASTQ files into subsequences from 12 specimens, each containing approximately 1.5 million unique sequences, required 2.3 hours and calculating k-mer enrichment required 7.7 minutes. The duration of these calculations scales approximately linearly with the number of specimens and sequences.

### K-TOPE algorithm

The K-TOPE algorithm (Code S1) is written in Python 3.6 (KTOPE.py). A usage guide for KTOPE is available (Text S1). First, there is a RAM-intensive step of loading k-mer enrichment data into memory as a dictionary. The enrichment dictionary for 250 specimens required approximately 4 GB of RAM. Then, a protein of interest is chosen for analysis and its sequence is loaded. This protein is tiled into k-mers of a set length. For this study, 5-mers were used. Each position in the protein sequence is assigned a frequency counter that starts at 0. The frequency counter of each sequence position contained in an enriched k-mer is incremented by the logarithm base 2 of the k-mer’s enrichment. For instance, if 3 k-mers that overlapped at a position had enrichments of 2, 4, and 8, the frequency for that position would be log_2_ 2 + log_2_ 4 + log_2_ 8 = 6. The frequency counters are compiled into a histogram which is smoothed using a moving window. For this analysis, the window had width 7 and used linear weighting with 1 in the center and 0.1 at the edges. Minima and maxima are identified in the smoothed histogram. All intervals between 2 minima that contain a maximum are used to define epitopes. Epitopes were limited to a minimum length of 6 and a maximum length of 15. Epitopes are scored using the area under the curve of the un-smoothed histogram. To assign statistical significance to each epitope, the epitope’s score is ranked in a list of scores for epitopes of the same length generated through an analysis of 10,000 random proteins. This rank is reported as a percentile in the distribution of random protein epitope scores. For this study, a percentile cutoff of 95% was used. For 12 specimens, analysis of 10,000 random proteins required 10.0 minutes.

After determining epitopes for individual specimens, K-TOPE can determine consensus epitopes for a population. Each epitope is characterized by a “centroid” which is the weighted central position of the epitope, indexed as a position in the protein sequence. Centroids for all epitopes that meet the percentile cutoff are compiled. They are then clustered using k-means to associate close centroids with the KMeans function from scitkit-learn [64]. A representative epitope is made for each cluster and kept if it meets a minimum prevalence in the population. Closely overlapping epitopes are removed and the final list is sorted by prevalence. Consensus epitopes can be determined for each protein in a proteome, generating a list of epitopes prevalent in a population. Determination of consensus epitopes for the *Rhinovirus A* genome polyprotein (P07210) for 250 specimens required 24.4 seconds. The proteome searches for viruses with human tropism, *Staphylococcus*, and *Streptococcus* for 250 specimens required 3.1, 2.3, and 1.9 hours, respectively.

We calculated expected membership of epitope groups by multiplying the proportions of the population that bound each epitope. For example, if epitope 1 was bound by 32% of the population and epitope 2 was bound by 67%, then the expected membership of epitope group ‘1+2’ would be 21%. We ranked the overlaps between K-TOPE derived epitopes and literature epitopes in a list of 10,000 randomly generated epitope overlaps to determine a p-value. To remove redundant epitopes found in the proteome searches, we used the PAM30 similarity matrix to align two epitopes and compare each position to calculate a similarity score. Epitopes that had similarity scores >10, were in the same protein, and were from different organisms were considered redundant. We removed the less prevalent of the two redundant epitopes.

The HSV analysis used “disease” group specimens to identify epitopes and “control” group specimens to subtract epitopes. Epitopes were identified in the disease group that met the epitope percentile cutoff (95%) and the minimum prevalence (30%). Then, all disease epitopes were evaluated in the control group. For an epitope to be considered disease-specific, its score had to be below the epitope percentile cutoff (80%) in all control specimens. To identify HSV2-specific epitopes that were also in the HSV1 proteome, we identified epitopes that exactly matched a subsequence in an HSV1 protein.

### Data visualization

Fig 1 was created using Inkscape. Histograms and heat maps were generated using the Matplotlib python module [65]. Bar graphs were generated using GraphPad Prism 7.

## Acknowledgements

The authors acknowledge the use of the Biological Nanostructures Laboratory within the California NanoSystems Institute, supported by the University of California, Santa Barbara and the University of California, Office of the President. We would like to acknowledge the work of Jack Reifert, Robert Pantazes, Chia-In Lin, Serra Elliott, and Kiho Song. We would like to acknowledge Linc Johnson for help with the initial conceptualization of this project.

## Supporting information

**S1 Fig. A comparison of histograms generated by K-TOPE when antibodies were added to serum or buffer.** Histograms were generated for antibodies against cMyc (P01106), V5 (P11207), and amyloid beta (P05067). The most prominent peaks were present regardless of whether antibodies were added to serum or buffer. This suggests that the binding signature of a single antibody was not obscured by the many other antibody specificities present in serum.

**S1 Table. The expected and actual membership of different epitope groups.** The expected membership of epitope groups was calculated by multiplying the proportions of the population that bound each epitope. For example, if epitope 1 was bound by 32% of the population and epitope 2 was bound by 67%, then the expected membership of epitope group ‘1+2’ would be 21%. Note that specimens in groups *only* bound the epitopes in the groups e.g. specimens in group ‘1’ did not bind ‘2’ or ‘3’. Generally, the actual and expected membership values agreed except for the ‘1+2+3’ group which had higher membership than expected and the ‘1+3’ group which had lower membership than expected (in bold).

**S2 Table. The average age for each epitope group.** The average age for the 138 specimens for which there was age data was 35. The ‘None’ group had an average age of 50 which was notably higher than the average age of 35 (in bold). Additionally, the ‘1+2+3’ group had a lower average age of 17 (in bold). This discrepancy suggests that older people targeted fewer *Rhinovirus A* epitopes.

**Table S3. Eight HSV2-specific epitopes were also in the HSV1 proteome.**

**Code S1. KTOPE software, written in Python 3.6.**

**Text S1. KTOPE usage guide.**

